# Analysis and Correction of Inappropriate Image Duplication: The *Molecular and Cellular Biology* Experience

**DOI:** 10.1101/354621

**Authors:** Elisabeth M. Bik, Ferric C. Fang, Amy L. Kullas, Roger J. Davis, Arturo Casadevall

## Abstract

The present study analyzed 960 papers published in *Molecular and Cellular Biology* (MCB) from 2009-2016 and found 59 (6.1%) to contain inappropriately duplicated images. The 59 instances of inappropriate image duplication led to 42 corrections, 5 retractions and 12 instances in which no action was taken. Our experience suggests that the majority of inappropriate image duplications result from errors during figure preparation that can be remedied by correction. Nevertheless, ~10% of papers with inappropriate image duplications in MCB were retracted. If this proportion is representative, then as many as 35,000 papers in the literature are candidates for retraction due to image duplication. The resolution of inappropriate image duplication concerns after publication required an average of 6 h of journal staff time per published paper. MCB instituted a pilot program to screen images of accepted papers prior to publication that identified 12 manuscripts (14.5% out of 83) with image concerns in two months. The screening and correction of papers before publication required an average of 30 min of staff time per problematic paper. Image screening can identify papers with problematic images prior to publication, reduces post-publication problems and requires significantly less staff time than the correction of problems after publication.

## Introduction

Recently we reported an analysis of 20,000 papers from 40 biomedical journals, published over a period of 20 years, in which approximately 1 in 25 papers contained at least one inappropriately duplicated image (1). The frequent occurrence of inappropriate image duplication in published papers is a major concern, because it reduces the integrity and credibility of the biomedical literature. At one end of the spectrum, inappropriate image duplications caused by simple errors in constructing figures raise concerns about the attention given to the preparation and analysis of data, while at the other end of the spectrum, problems resulting from deliberate image manipulation and fabrication indicate misconduct. Increased awareness of such image duplications has resulted from post-publication peer review websites such as PubPeer and discussions on social media (2). Whereas simple errors found in published studies can be addressed by a correction, deliberate image manipulation or fabrication can lead to retraction of a paper (3).

Inappropriate image duplications undermine the quality of the literature and can necessitate a considerable investment of time and resources by authors and journals when discovered after publication of a scientific paper. However, we presently lack information on the causes for the inappropriate image duplications, since neither cause nor intent can be reliably inferred from inspecting images in published articles. We categorized inappropriate image duplications as simple duplications (category 1), shifted duplications (category 2) or duplications with alterations (category 3), with category 1 most likely to result from honest error, while categories 2 and 3 have an increased likelihood of resulting from outright falsification or fabrication. A follow-up analysis of a subset of these papers found that several variables including academic culture, peer control, cash-based publication incentives and national misconduct policies were significantly associated with duplications in categories 2 and 3, suggesting that these variables might affect scientific integrity (4). In the present study, we sought to determine whether an investment by a journal to scan images in accepted manuscripts prior to publication could resolve image concerns in less time that was required to address these issues after publication.

The mission of the journals published by the American Society for Microbiology (ASM) is to publish high-quality scientific articles that have been rigorously peer reviewed by experts and evaluated by academic editors (5). In 2013, the ASM journal *Molecular and Cellular Biology* (MCB) instituted a program to analyze the figures in all accepted manuscripts before publication (6), modeled after a similar program used by the *Journal of Cell Biology* (7, 8). In this study, we applied the approach used previously (1) to published papers in the journal MCB, and followed up the findings with a process that included contacting the authors of the papers. Consequently, we are now able to provide information as to how inappropriate image duplications occur. In addition, a set of manuscripts accepted for publication in MCB was inspected prior to publication for spliced, beautified, or duplicated images. For both sets of papers, the time and effort spent on following up on these papers was recorded. The results provide new insights into the prevalence, scope and seriousness of the problem of inappropriate image duplication in the biomedical literature.

## Methods

### Published papers set

Papers published in 2009-2016 in MCB were inspected visually for inappropriate image duplication. For each year, issues 1-12 (January-June) were selected, and the first 10 papers in each issue containing photographic images were screened. Thus, 120 papers were inspected per publication year, resulting in a total of 960 papers screened. Since almost all MCB papers contain photographic images, no specific search term was used, but papers were only counted if they contained photographic images.

### Image inspection

Published papers were scanned using the same procedure used in our prior study (1). Briefly, one person (EMB) scanned published papers by eye for image duplications in any photographic images or FACS plots. Problematic images were also inspected by two additional authors (AC and FCF). Such duplicated images fell into three general categories: simple duplications, duplications with repositioning, and duplications with alterations (1). As in the previous study (1), cuts and beautifications were not scored as problematic. EMB was not aware of the year in which MCB started increased screening (see below) for image problems while she screened journals. The image allegations were confirmed using ORI forensic software by the MCB Production Department. Decisions as to whether to pursue the allegations by contacting authors were based on this analysis. Each published paper containing suspected image duplication problems was reported to the Editor-in-Chief of MCB. The EIC then requested clarification from the corresponding author(s) regarding concerns with the figure using the category classification described above. The EIC followed up on all concerns from 2010 and on potential concerns in Categories 2 & 3. Category 1 concerns were handled by ASM staff.

### Prospective screening of manuscripts before publication

Starting in January 2013, all MCB manuscripts accepted for publication were screened for image duplications and other problems, including undisclosed cuts and beautifications (which were not counted in the screen of the published papers described above). For this study, the time to inspect these figures in manuscripts accepted from January 14, 2013 to March 21, 2013 was recorded. In the case of image problems, the authors were contacted and asked to explain and/or remake the figure. Corrections and retractions followed COPE guidelines (https://publicationethics.org/resources/guidelines).

## Results

### Inappropriate duplications in MCB published papers

A set of 960 papers published in MBC between 2009 and 2016, including 120 randomly selected papers per year, was screened for image duplication. Of these, 59 (6.1%) papers were found to contain inappropriately duplicated images. The distribution of these showed a decline since 2013, when the screening of accepted manuscripts was introduced (Figure 1). From 2009-2012, the average percentage of image duplication was 7.08%, while after the introduction of screening accepted manuscripts in 2013, the percentage was 3.96%, a significant decrease (t test; p<0.01).

**Figure 1.**
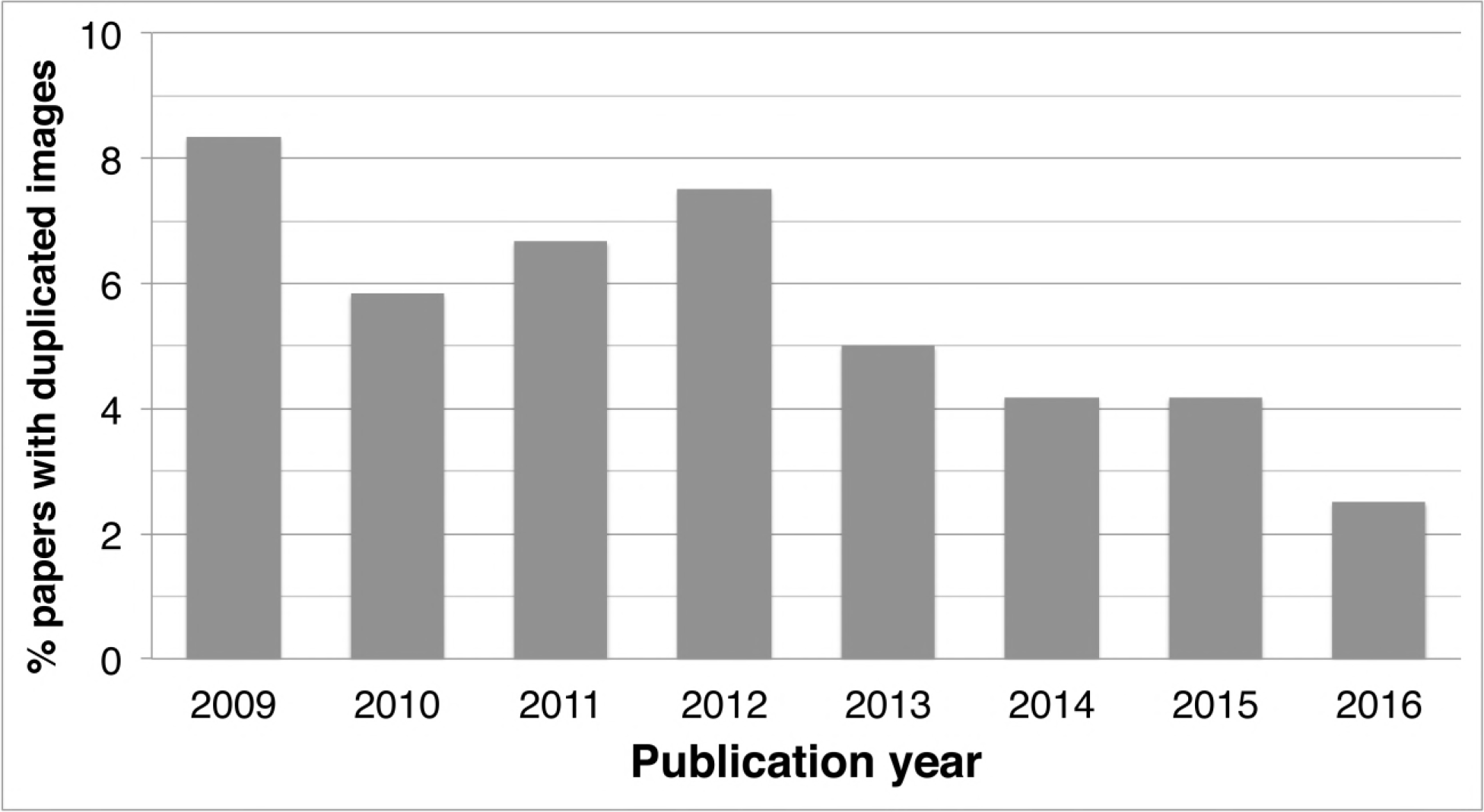
Percentage of papers published in ASM’s *Molecular and Cellular Biology* containing duplicated images. Inspection of manuscripts pre-publication started in 2013.

### Investigation by ASM staff into published papers with image duplication

The 59 papers with inappropriate image duplications in MCB were investigated by contacting the corresponding authors and requesting an explanation for the apparent problem. The 59 instances of inappropriate image duplications led to 42 corrections, 5 retractions and 12 instances in which no action was taken (Table 1). The reasons for not taking action included origin from laboratories that had closed (2 papers), resolution of the issue in correspondence (4 papers), and occurrence of the event more than six years earlier (6 papers), consistent with ASM policy and Federal regulations established in 42 CFR § 93.105 for pursuing allegations of research misconduct. Of the retracted papers, one contained multiple image issues such that a correction was not an appropriate remedy, and for another retracted paper, the original and underlying data was not available, but the study was sufficiently sound to allow resubmission of a new paper for consideration, which was subsequently published.

**Table 1.** Summary of results and comparison of image duplication problems in published MCB papers and accepted MCB manuscripts.

| Action | Explanation | Papers with Image duplication problems |  |
| --- | --- | --- | --- |
|  |  | Post publication (59, 6.1%)<sup>1</sup> | Pre-publication (12, 14.5%)<sup>2</sup> |
| None | Duplication could not be confirmed or not a strong case | 3 (5.1%) | 0 |
|  | Duplication was legitimate | 1 (1.7%) | 0 (0%) |
|  | Lab had closed after submission of paper, not pursued | 2 (3.3%) | 0 (0%) |
|  | Older than 6 years | 4 (6.8%) | 0 (0%) |
|  | Authors did not reply | 2 (3.3%) | 0 (0%) |
|  | Authors provided original blot showing no duplication | 1 (1.7%) | 0 (0%) |
| Correction | Simple duplication during figure assembly | 40 (67.8%) | 11 |
|  | Error in figure assembly | 1 (1.7%) | 1<sup>4</sup> |
| Retraction (or rejection of manuscript) | Too many errors for simple correction | 3 (5.1%) | 0 |
|  | Intention to mislead suspected | 2 (3.3%) | 0 |
| Staff effort | Emails sent to resolve problems | 1400-1600<sup>3</sup> | N.A.<sup>5</sup> |
|  | Average time spent per paper | 6 h | 1.5 h |

<sup>1</sup> Published papers (n=960), 2009-2016
<sup>2</sup> Accepted papers (n=83), 2 months in 2013
<sup>3</sup> Email estimate includes EIC correspondence.
<sup>4</sup> Analysis revealed potential case of non-uniform enhancements
<sup>5</sup> N.A. means not available

### Analysis of inappropriate image duplications

Authors who were contacted about image irregularities most frequently reported errors during assembly of the figures. The most commonly reported error was the accidental inclusion of the same blot or image twice. Other commonly reported mistakes were the selection of the wrong photograph, the assembly of figure panels with mock photographs that were not properly replaced, etc.

### Time effort for published papers

For the 59 papers published with potential image duplication concerns, the ASM publication staff members recorded ~580 emails pertaining to these cases, or an average of ~10 emails per case (range 4-103). In addition, at least two phone conversations with authors took place, each approximately 1 h. The Production Editor and Assistant Production Editor handled ~800 emails in their folders regarding these corrections. In addition, for 20 papers the Editor in Chief (EIC) was involved in communications with the authors, which involved a total of 244 emails (range per paper 4-29) or an average of 12.2 messages per paper. Including the EIC time would add another 61 h (~15 min × 244 emails). The exact content of these emails was not disclosed to any individuals outside of the MCB ethics panel. The breakdown of the Production Staff emails were: correspondences with staff members to keep them apprised of what had been received, discussions about wording (since each item needed individual assessment of the appropriate approach), or logistical details regarding retracted or republished papers. Correspondences with authors comprised the next largest category (less than half the amount of staff correspondence), followed by correspondences with the EIC. Correspondences with the printer was the smallest category. Hence, the problem of inappropriate image duplication after publication imposed a large time burden on the journal, with an average of 6 h of combined staff time (1400 emails estimated to take 15 min each to write and follow up per 59 papers) spent to investigate and follow-up each paper.

### Screening of manuscripts prior to publication

Analyzing the papers with inappropriately duplicated images as a function of time revealed a decline in incidence beginning in 2013, which coincided with a change in the editorial process to include pre-publication screening for image problems (Figure 1). During a period of 2 months in the beginning of 2013, 83 papers were accepted with 452 images inspected. In this recording period, 12 papers (14.5%) were detected in which an image concern (duplication or undisclosed cuts) was identified. The percentage of papers flagged during pre-publication screening was higher than the frequency of duplicated images detected in published papers, because beautification or undisclosed cuts were flagged as well. Prior to this time, no manuscript was rejected by MCB because of image duplication, but starting in 2013, after the introduction of pre-publication screening, the percentage of manuscripts rejected for image problems steadily increased (Figure 2).

**Figure 2.**
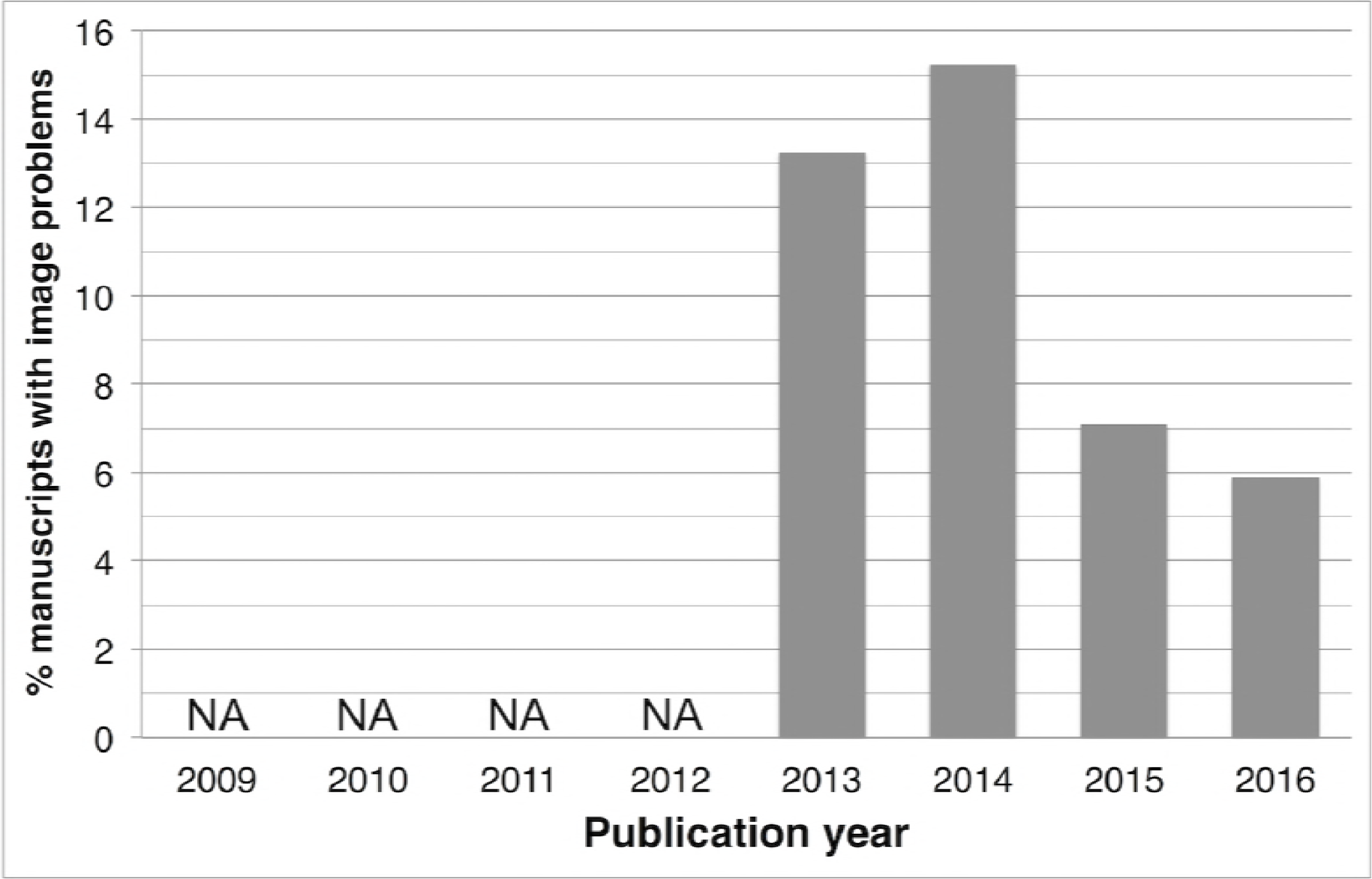
Percentage of accepted MCB manuscripts that were found to have problematic images, 2013-2016. No screening for problematic images was done before 2013 and NA means non-applicable.

### Outcome of pre-publication screening of manuscripts

During the recording period in 2013, 84 manuscripts were screened, and 12 manuscripts were flagged for containing duplications or other irregularities. The authors of each manuscript were contacted for follow-up by the handling editor. In 11 cases, the problem could be corrected by the submission of a new version of the figure, while in 1 instance, the authors provided the original data to show that the figure did not misrepresent the original data.

### Time effort for manuscripts

When image screening was first instituted by MCB in 2013, time records were maintained for approximately two months to ascertain the time cost of this procedure. The total time required to inspect all images in the 84 manuscripts screened during this period was 687 minutes (8.3 minutes per screened paper). The total time required for reporting and following up of ethical concerns found in 12 papers was 375 minutes of ASM staff time, not counting the time devoted by the EIC to addressing these problems. Thus, the time effort of ASM staff/editors translated into 31.3 minutes per manuscript.

## Discussion

Here we report the first detailed investigation of inappropriate image duplications in biomedical research papers and a systematic process for their correction. By focusing on one journal within the ASM journals portfolio, we were able to determine the outcome of image concerns. The most reassuring outcome of our findings is that the majority of inappropriate image duplications resulted from errors during figure construction that could be easily corrected by the authors. The finding that 5.5% of MCB articles had inappropriate image duplications is a percentage consistent with prior findings involving over 40 journals (1). This confirmation is noteworthy because the approach used in the current study differs from prior work in that it focused on a single journal with a 120-paper sample for each of six publication years. Of concern is that approximately 10% of the papers containing problematic images required retractions after the adjudication process, due to apparent misconduct, an inadequate author response, or errors too numerous for an authors’ correction. Other efforts to investigate causes of inappropriate image duplication for papers published at two other American Society of Microbiology journals, *Journal of Virology* and *Infection and Immunity*, including some from a prior study (1), produced retraction rates ranging from 2.9 (1 of 35) to 21% (4 of 19), respectively, which yields an average of 10.6 ± 8.1% for the three journals.

Research misconduct has always existed, but this topic has been of increasing concern in recent years in view of several high profile scandals, a perceived reproducibility crisis and an epidemic of retracted papers, most of which are due to misconduct (9). The actual number of compromised papers in the extant literature is unknown, but our observations permit some estimates. Although extrapolation from three American Society of Microbiology journals to the general biomedical literature must be made with caution, our study allows a rough estimate of the number of seriously compromised papers in print. Based on the average percent retraction from the three journals, the 95% confidence interval ranges from 1.5-19.8%. If 3.8% of the 8,778,928 biomedical publications indexed in PubMed from 2009-2016 (http://dan.corlan.net/medline-trend.html) contain a problematic image (1), and 10.6% (CI 1.5-19.8%) of that group contain images of sufficient concern to warrant retraction, then we can estimate that approximately 35,000 (CI 6,584-86,911) papers are candidates for retraction due to image duplication. These numbers are almost certainly an overestimate since not all papers in the literature have images of the type studied here. On the other hand, we only screened for visible duplications, and papers might contain additional problems in graphs, tables, or other datasets that are less easy to find, suggesting that this could also be an underestimate. Whatever the actual number, it is clear that the number of compromised papers in the literature is significant. The continued presence of compromised papers in the literature could exert pernicious effects on the progress of science by misleading investigators in their fields. Nevertheless, even the most liberal estimates the total number of papers that are candidates for retraction represent a very small percentage of the literature. Our findings are consistent with other studies reporting that a significant number of papers in the literature have problems associated with misconduct (10, 11).

Our study also documents the potential value of increased journal vigilance for reducing inappropriate image duplications in published papers. A significant reduction in the number of inappropriate image identified in MCB papers was observed after initiation of dedicated image inspections by the journal in 2013 (6). Increased vigilance reduces problematic images by identifying and correcting errors before publication and by heightening awareness among authors to prevent such problems. However, such efforts come at considerable time and financial costs to the journal. The time invested in inspecting manuscripts pre-publication was approximately 8.3 minutes per paper, and the identification of a problematic image resulted in additional time investment in communicating with authors and deciding if a problem raised an ethical concern. Additional costs to science include the time taken by the authors to correct figures and the delays in publication. However, these costs may be significantly lower than the overall cost associated with discovery of image duplication after publication, which triggers an investigation by the journal that consumes considerable time, as is evident from the average of 10 emails per case, to outcomes including publication of corrections and retractions. In our analysis, we found that following up on problematic images before publication costs about 30 min per problematic paper, whereas the time spent to follow up similar issues after publication, not including EIC time, was 6 h per paper, which is twelve times greater. Hence, even though the majority of inappropriate image duplications result from simple errors in assembling figures, their occurrence once identified imposes considerable costs to journals and authors, and by extension, to the scientific enterprise. Identifying image problems before publication, even though this requires additional time for journal staff, might save journals time in the end by preventing problematic images from appearing in published papers. In addition, identifying potential problems before publication protects authors’ reputations and prevents the collateral damage to the reputations of all authors of a retracted paper (12).

Peer review is a cornerstone of science (13, 14), which is primarily designed to look for fundamental errors in experimental setup and data analysis. Most peer reviewers do not have the expertise to analyze papers for scientific misconduct. Consequently, the responsibility of screening for plagiarism, falsification, fabrication, and other forms of science misconduct often lies with editors (15). Although sloppiness and misconduct have always existed in science, the problem may be becoming more acute because of advances associated with the information revolution. The ability to cut-and-paste text or images combined with availability of software to manipulate and generate photographic images gives authors powerful tools that can be misused. Our prior study noted that the problem of inappropriate image duplications was largely a 21^st^ century phenomenon temporally associated with the proliferation of software for image construction (1). However, the information revolution has also provided tools to reduce error and abuse. Some publishers, including ASM, already perform routine screening of manuscripts using plagiarism-detection software. Combined with manual curation and supervision, these tools work reasonably well (11, 16). However, identifying image duplication of the types reported here and in our prior study (1) is more challenging and dependent on individuals capable of spotting suspicious patterns. We noted that the pre-screening process at MCB is quite good at picking up spliced images but poor at finding image duplications of the type reported in this study. Hence, without routine screening by individuals who are gifted at identifying image duplications and modifications, it is likely that the type of image problems identified here will continue (1). Although detecting image problems is difficult, the recent development of improved software tools appears promising (17).

The finding that most inappropriate image duplications result from sloppiness and error during figure construction but impose large costs to authors and journals for their correction indicates that greater efforts to prevent such errors should be instituted by research laboratories. Given that most figures are currently constructed by authors themselves, it may be possible to reduce the prevalence of image problems by asking others in the laboratory that are not directly involved with the current research to participate in figure construction or review. Prior to the availability of image editing software, figures for research papers were usually made by individuals who specialized in this activity and were not involved in data collection. We note that in our previous study we found no instances of inappropriate image duplication prior to 1997 (1). We hypothesize that prior to the availability of software that allowed authors to construct their own figures, the discussions between photographers or illustrators and authors combined with the separation of data generation from figure preparation reduced the likelihood of these types of problems.

In addition, providing clear guidelines for the preparation of photographic images as part of a journal’s instructions for authors is helpful. For example, instructions might include rules about how to disclose cuts in Western blots, the requirement of each experiment to have their own control (e.g. β-actin or globin) protein control blots (no re-use of these blots allowed), etc. Examples of such guidelines currently exist (18). ASM maintains an ethics portal in its website with information that may be helpful to authors (http://journals.asm.org/site/misc/ethicsportal.xhtml)

In summary, we confirmed our prior results by inspecting a single journal using a systematic approach and provide insights into the causes of inappropriate image duplication in research papers. The results provide both reassurance and concern regarding the state of the biomedical literature. We are reassured that the majority of duplication events result from errors that do not compromise validity of the scientific publication and are amenable to correction, notwithstanding the cost of considerable time investment on the part of the journal staff, editors and authors. However, of concern is the significant minority of papers with inappropriate image duplications that result in retractions, suggesting that the current biomedical research literature contains many such publications that warrant retraction. At the very least, our findings suggest the need for both authors and journals to redouble their efforts prevent inappropriate image duplications.

## REFERENCES

1. Bik EM, Casadevall A, & Fang FC (2016) The Prevalence of Inappropriate Image Duplication in Biomedical Research Publications. MBio 7(3).

2. Knoepfler P (2015) Reviewing post-publication peer review. Trends Genet 31(5):221–223.

3. Fang FC & Casadevall A (2011) Retracted science and the retraction index. Infect. Immun 79(10):3855–3859.

4. Fanelli D, Costas R, Fang FC, Casadevall A, & Bik EM (2018) Testing Hypotheses on Risk Factors for Scientific Misconduct via Matched-Control Analysis of Papers Containing Problematic Image Duplications. Science and engineering ethics.

5. Casadevall A, et al. (2016) ASM Journals Eliminate Impact Factor Information from Journal Websites. MBio 7(4).

6. Kullas AL & Davis RJ (2017) Setting the (Scientific) Record Straight: Molecular and Cellular Biology Responds to Postpublication Review. Molecular and cellular biology 37(11).

7. Rossner M (2002) Figure manipulation. assessing what is acceptable 158(7):1151–1151.

8. Yamada KM & Hall A (2015) Reproducibility and cell biology. The Journal of cell biology 209(2):191–193.

9. Fang FC, Steen RG, & Casadevall A (2012) Misconduct accounts for the majority of retracted scientific publications. Proc. Natl. Acad. Sci. U. S. A 109(42):17028–17033.

10. Looi LM, Wong LX, & Koh CC (2015) Scientific misconduct encountered by APAME journals: an online survey. The Malaysian journal of pathology 37(3):213–218.

11. Taylor DB (2017) JOURNAL CLUB: Plagiarism in Manuscripts Submitted to the AJR: Development of an Optimal Screening Algorithm and Management Pathways. AJR. American journal of roentgenology 208(4):712–720.

12. Bonetta L (2006) The aftermath of scientific fraud. Cell 124(5):873–875.

13. Glonti K, et al. (2017) A scoping review protocol on the roles and tasks of peer reviewers in the manuscript review process in biomedical journals. BMJ open 7(10):e017468.

14. Kronick DA (1990) Peer review in 18th-century scientific journalism. Jama 263(10):1321–1322.

15. Babalola O, Grant-Kels JM, & Parish LC (2012) Ethical dilemmas in journal publication. Clinics in dermatology 30(2):231–236.

16. Lykkesfeldt J (2016) Strategies for Using Plagiarism Software in the Screening of Incoming Journal Manuscripts: Recommendations Based on a Recent Literature Survey. Basic & clinical pharmacology & toxicology 119(2):161–164.

17. Butler D (2018) Researchers have finally created a tool to spot duplicated images across thousands of papers. Nature 555(7694):18.

18. Collins S, Gemayel R, & Chenette EJ (2017) Avoiding common pitfalls of manuscript and figure preparation. Febs j 284(9):1262–1266.

